# Loss of consumers constrains phenotypic evolution in the resulting food web

**DOI:** 10.1101/527416

**Authors:** Matthew A. Barbour, Christopher J. Greyson-Gaito, Arezoo Sootodeh, Brendan Locke, Jordi Bascompte

## Abstract

The loss of biodiversity is altering the structure of ecological networks; however, we are currently in a poor position to predict how these altered communities will affect the evolutionary potential of remaining populations. Theory on adaptive landscapes provides a framework for predicting how selection constrains phenotypic evolution, but often treats the network context of evolving populations as a “black box”. Here, we integrate ecological networks and adaptive landscapes to examine how changes in food-web structure shape evolutionary constraints. We conducted a field experiment that removed a guild of larval parasitoids that imposed direct and indirect selection pressures on an insect herbivore. We then measured herbivore survival as a function of three key phenotypic traits to estimate directional, quadratic, and correlational selection gradients in each treatment. We used these selection gradients to characterize the slope and curvature of the adaptive landscape to understand the direct and indirect effects of consumer loss on evolutionary constraints. We found that the number of traits under directional selection increased with the removal of larval parasitoids, indicating greater selective constraints on the trajectory of evolutionary change. Similarly, we found that the removal of larval parasitoids altered the curvature of the adaptive landscape in such a way that tended to decrease the evolvability of the traits we measured in the next generation. Our results suggest that the loss of trophic interactions can impose greater constraints on phenotypic evolution. This indicates that the simplification of ecological communities may constrain the adaptive potential of remaining populations to future environmental change.

**Impact Summary:** The loss of biodiversity is rewiring the web of life; however, it is uncertain how this will affect the ability of remaining populations to evolve and adapt to future environments. To gain insight to these effects, we conducted a field experiment that either maintained a natural community of predators or removed all but one of the predators that was able to impose selection on a common prey. We found that the loss of predators acted to constrain prey evolution toward a particular combination of traits. Moreover, we found that the loss of predators could make it more difficult for prey to adapt to uncertain future environments. Taken together, our results suggest that the simplification of the web of life may constrain the adaptive potential of remaining populations.

## Introduction

The adaptive landscape provides a powerful framework for understanding how natural selection has shaped the evolution of biodiversity —from genes to phenotypes to species (Wright, 1931; Simpson, 1944; Arnold et al., 2001). More than a metaphor, the adaptive landscape links quantitative genetic and phenotypic variation to evolution by natural selection (Lande, 1979; Arnold and Wade, 1984*a*,*b*), giving insight to the direct and indirect effects of selective constraints (Arnold, 1992). Ecological interactions often play a key role in shaping selective constraints, as evidenced by the role of antagonistic and mutualistic interactions in driving evolutionary change (Schluter, 2000; Abrams, 2000; Bronstein et al., 2006). Although there is clear evidence that species interactions can shape the adaptive landscape, we still have a poor understanding of how the adaptive landscape is shaped by a community of interacting species (McPeek, 2017; terHorst et al., 2018). Resolution on how change in ecological communities shape phenotypic evolution is urgently needed though, given the rapid losses of biodiversity we are observing in the Anthropocene (Scheffers et al., 2016).

Ecological networks, such as food webs describing who-eats-whom, provide an explicit representation of the direct and indirect effects that emerge in a community of interacting species (Bascompte and Jordano, 2014; McCann, 2012). Here, we integrate ecological networks and adaptive landscapes to understand how ecological communities constrain evolutionary change. The direct and indirect effects of natural selection can be inferred by quantifying the slope and curvature of the adaptive landscape (Arnold, 1992). For example, the slope is determined by directional selection gradients acting on each phenotypic trait and influences the trajectory of evolutionary change (Lande, 1979; Arnold, 1992). Selective constraints on evolution increase with the number of traits under directional selection, as this diminishes the number of optimal solutions (Arnold, 2003). The curvature of the adaptive landscape is governed by nonlinear selection gradients and acts to indirectly constrain evolution through its effect on genetic constraints (Arnold, 1992; Hansen and Houle, 2008). Genetic constraints are largely governed by a population’s G-matrix —the additive genetic variances and covariances between traits (Hansen and Houle, 2008). Stabilizing selection acts to erode genetic variance in traits, which can impose a constraint on the ability of those traits to respond to future selection (Hansen and Houle, 2008). Correlational selection may alter the genetic covariance between traits, which may facilitate or hinder future adaptation depending on the pattern of selection on those traits (Hansen and Houle, 2008). If we want to predict how ecological communities shape phenotypic evolution, we must understand how ecological networks shape the adaptive landscape.

The loss of biodiversity is altering the structure of ecological networks, which may influence evolutionary constraints in a number of ways. For example, in a food web, if different consumers impose directional selection on different traits of a shared resource, then a more diverse consumer community may constrain evolution by increasing the number of traits under selection. Alternatively, if consumers impose selection on different values of a trait, then their selective effects would cancel each other out in more diverse communities. To examine these different possibilities, we conducted a field experiment that removed a consumer guild that directly and indirectly impacts fitness of an abundant insect herbivore (*Iteomyia salicisverruca*, Family Cecidomyiidae; fig. 1). The larvae of this herbivore induce tooth-shaped galls when they feed on the developing leaves of willow trees (*Salix* sp., Russo, 2006). These galls protect larva from attack by generalist predators (e.g. ants, spiders), but they suffer high mortality from egg and larval parasitoids (Barbour et al., 2016). We manipulated food-web structure by either removing larval parasitoids (removal food web) or allowing both egg and larval parasitoids to impose selection on gall midge traits (original food web, fig. 1). The relative abundance of these larval parasitoids is much lower than the egg parasitoid (Barbour et al., 2016), and thus represents a plausible extinction scenario for this system. Larval parasitoids also impose indirect effects on gall midge fitness through intraguild predation on the egg parasitoid (fig. 1). We applied modern statistical methods to quantify how changes in food-web structure altered the slope and curvature of the gall midge’s adaptive landscape. Taken together, our study gives insight to how the loss of biodiversity may constrain the evolution of interacting populations.

**Figure 1:**
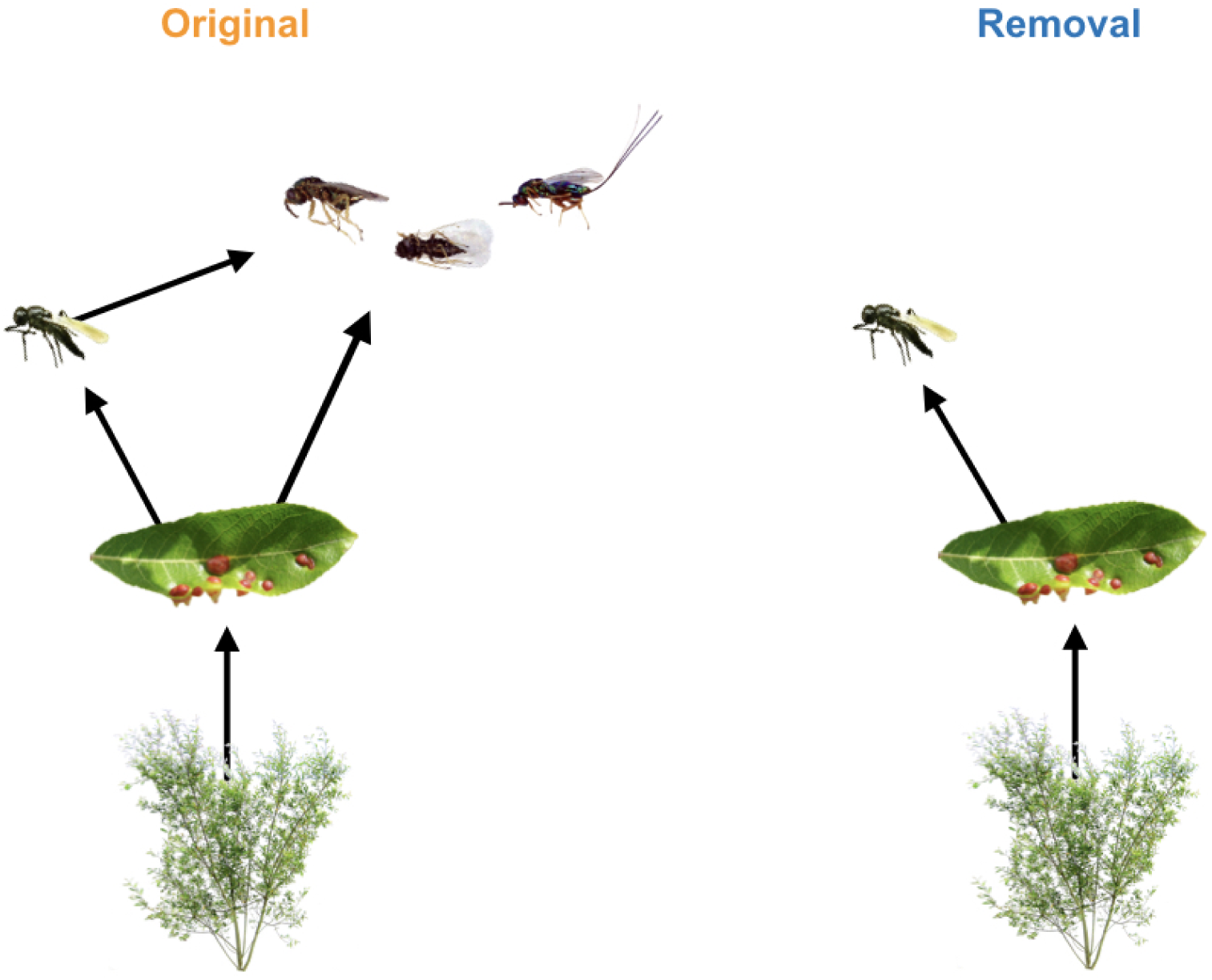
Experimental manipulation of food-web structure associated with a leaf-galling midge (*Iteomyia salicisverruca*) feeding on the willow *Salix hookeriana*. Black arrows denote the flow of energy in this network of trophic interactions. In the original food web, we allowed the full suite of egg and larval parasitoids to impose selection. In the removal food web, we used mesh bags to exclude the guild of larval parasitoids, only allowing the egg parasitoid (*Platygaster* sp.) to impose selection. Note that larval parasitoids also impose indirect effects on gall midge fitness through intraguild predation on the egg parasitoid. Larval parasitoids include the following species (from left to right): *Mesopolobus* sp. (Family: Pteromalidae); *Tetrastichus* sp. (Family: Eulophidae); and *Torymus* sp. (Family: Torymidae).

## Methods

### Study Site

We conducted our study within a four-year old common garden experiment of coastal willow (*Salix hookeriana*) located at Humboldt Bay National Wildlife Refuge (HBNWR) (40 40’53“N, 124 12’4”W) near Loleta, California, USA. This common garden consists of 26 different willow genotypes that were collected from a single population of willows growing around Humboldt Bay. Stem cuttings of each genotype (25 replicates per genotype) were planted in a completely randomized design in two hectares of a former cattle pasture at HBNWR. Willows at our study site begin flowering in February and reach their peak growth in early August. During this study, willows had reached 5 - 9m in height. Further details on the genotyping and planting of the common garden are available in Barbour et al. (2015).

### Manipulating Food-web Structure

We setup our food-web manipulation across 128 plants soon after galls began developing on willows in early June of 2013. These 128 plants came from eight different plant genotypes that spanned the range of trait variation observed in this willow population (Barbour et al., 2015). For the original food web (eight replicates per genotype), we used flagging tape to mark 14 galled leaves per plant (~30 larvae), allowing the full suite of egg and larval parasitoids to impose selection. Marking galls with flagging tape ensured that all galls had similar phenology when we collected galls later in the season. For the removal food web, we enclosed 14 galled leaves with 10×15cm organza bags (ULINE, Pleasant Prairie, WI, USA) to exclude three parasitoid species that attack during larval development. This treatment did not exclude the egg parasitoid *Platygaster* sp., which attacks prior to gall initiation (larva initiate gall development in Cecidomyiid midges: Gagné, 1989). It was not possible to reciprocally manipulate parasitoid attack (i.e. exclude egg parasitoid, but allow larval parasitoids), because it was not possible to identify midge oviposition sites prior to gall formation. In late August, we collected marked and bagged galls from each plant, placed them into 30 mL vials and kept them in the lab for 4 months at room temperature. We then opened galls under a dissecting scope and determined whether larvae survived to pupation (our measure of fitness) or were parasitized. We did not include other sources of mortality in our analyses, such as early larval death, as they could influence the expression of the gall phenotype and confound estimates of selection. For the food-web treatment that excluded larval parasitoids, we further restricted our data by removing any incidental instances of parasitism by a larval parasitoid. This represented less than 3% of the observations in this food-web treatment and allowed us to focus our inferences of selection on those imposed by the egg parasitoid. Our final dataset contains survival estimates for 1285 larvae from 613 galls and 111 plants.

### Measuring Phenotypic Traits

We collected data on three different traits that we expected to influence larval survival based on previous work in this system (Barbour et al., 2016) and other work with gall midges (Weis et al., 1983; Heath et al., 2018). First, we measured gall diameter as the size of each gall chamber to the nearest 0.01 mm at its maximum diameter (perpendicular to the direction of plant tissue growth). Previous work in this system has shown that larger galls are associated with higher survival (Barbour et al., 2016). Second, we measured clutch size by counting the number of chambers in each gall (Weis et al., 1983; Heath et al., 2018). All larvae collected from the same multi-chambered gall were scored with the same clutch size. Third, we measured oviposition preference as the density of larvae observed on a plant in an independent survey. We did this by randomly sampling five branches per tree and counting the number of individual gall chambers (number of larvae). We then converted these counts to a measure of larval density per 100 shoots by counting the number of shoots on the last branch we sampled. All larvae collected from the same plant were scored with the same oviposition preference. Measuring larval densities on plants in the field is a common method for measuring oviposition preference (Gripenberg et al., 2010); however, caution must be taken in inferring ‘preference’ as larval densities can be influenced by processes other than preference (Singer, 1986). Fortunately, a couple features of our study system suggest that larval densities may be a good proxy for oviposition preference. For example, since our data comes from a randomized placement of plant genotypes in a common garden, there is no consistent bias in which plant genotypes females are exposed to while searching for oviposition sites. Also, egg predation is a minor source of mortality for galling insects in general (Hawkins et al., 1997); therefore, we do not expect any prior egg predation to bias our estimates of observed larval densities.

### Quantifying the Adaptive Landscape

Inferring adaptive landscapes assumes that trait distributions are multivariate normal (Lande and Arnold, 1983). To approximate this assumption, we log-transformed clutch size and square-root transformed oviposition preference. We then scaled all phenotypic traits (mean=0 and SD=1) across treatments prior to our analyses (detailed below) to ensure that our estimates of selection were comparable across traits and with other studies.

Our analysis consisted of four parts. First, we used generalized linear mixed models (GLMM) to quantify selection surfaces —linear and nonlinear relationships between absolute fitness (*W*) and standardized phenotypic traits (*i*) of individuals —in each food-web treatment. Second, we translated selection surfaces into the scale of relative fitness (*w*) in order to calculate standardized selection gradients. Third, we used our estimates of directional selection gradients to characterize the slope of the adaptive landscape, which we used to measure selective constraints. Finally, we estimated the curvature of the adaptive landscape and used a simulation to explore its effects on the adaptive potential of the gall midge population in the next generation.

#### Selection surfaces

Since larval survival was our measure of absolute fitness, we used a GLMM that assumed a binomial error distribution (and logit-link function). To approximate the selection surface, we modelled larval survival as a function of food-web treatment as well as linear (*α*_*i*_), quadratic (*α*_*ii*_), and linear interactions (*α_ij_*) between each trait. We also allowed these selection surfaces (*α*) to vary between food-web treatments. Note that to obtain valid estimates of linear selection surfaces, we removed nonlinear terms prior to estimating linear relationships (Lande and Arnold, 1983). Other approaches have been advocated for approximating selection surfaces (Schluter, 1988); however, our approach enables us to calculate selection gradients, and thus is more appropriate for approximating the adaptive landscape (Arnold, 2003). To account for the nonindependence of clutch size (measured at gall level) and oviposition preference (measured at plant level) as well as any independent effects of willow genotype on larval survival, we modelled gall ID nested within plant ID nested within genotype ID as random effects. Although statistical models with random effects are not common in analyses of natural selection, we think that modelling random effects can mitigate biased estimates of selection due to environmental covariances between traits and fitness (Rausher, 1992). Since our end goal was to characterize the relationship between mean trait values and mean fitness (adaptive landscape), we assumed the mean value of our random effects (i.e. setting them to zero) when calculating selection surfaces. We then used parametric bootstrapping (1,000 replicates) to estimate the effect of food-web treatment on larval survival as well as selection surfaces in each food-web treatment. To determine whether trait-fitness relationships differed between food-web treatments, we calculated the difference in bootstrapped replicates between treatments.

#### Selection gradients

Selection gradients cannot be estimated directly from GLMMs of selection surfaces because the response is in terms of absolute fitness and the coefficients are on a nonlinear scale. For example, the coefficients in the above model measure the change in the logarithm of the odds of surviving (i.e. ln{*W*(*z*)/[1 − *W*(*z*)]}) associated with 1 SD change in a trait with all other traits held fixed at their mean. Therefore, we used the method developed by Janzen and Stern (1998) to translate selection surfaces from the above model into the scale of relative fitness in order to calculate directional (*β*_*i*_), quadratic (*γ*_*ii*_), and correlational (*γ*_*ij*_) selection gradients. Briefly, this method calculates the average gradient of selection surfaces by multiplying the average of *W*(*z*)[1 − *W*(*z*)] by each regression coefficient (e.g. *α*_*i*_, *α*_*ii*_, or *α_ij_*). We then divided this average gradient by the mean fitness 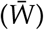 to put it on the scale of relative fitness, and thus interpretable as a selection gradient. We estimated selection gradients separately for each food-web treatment. We also determined whether selection gradients differed between food-web treatments by calculating the difference in bootstrapped replicates between treatments. Note that we doubled all quadratic terms prior to calculating selection gradients to put them on the same scale as estimates of directional and correlational selection (Stinchcombe et al., 2008).

#### Selective constraints

The number of selective constraints is determined by the slope of the adaptive landscape, which in our study corresponds to:

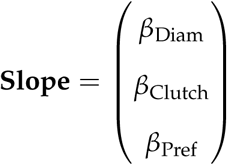

where each *β*_*i*_ corresponds to the directional selection gradient acting on each trait. By comparing the number of directional selection gradients that show clear evidence of contributing to the slope (i.e. 95% CI does not overlap zero) in our food-web treatments, we can infer the effect of food-web structure on selective constraints. This gives insight as to whether there is a specific combination of phenotypes favored by natural selection (i.e. more traits under directional selection), or whether multiple trait combinations have comparable fitness (i.e. less traits under directional selection).

#### Genetic constraints

The indirect effects of selection on the G-matrix (∆G = G(*γ* − *ββ*^T^)G) are governed by the curvature of the adaptive landscape (C = *γ* − *ββ*^T^), which in our study corresponds to:

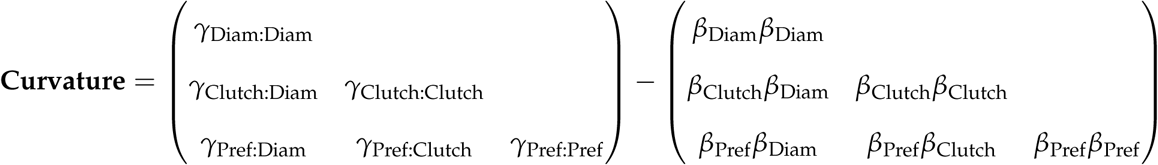

where each *γ*_*ii*_ (diagonal) corresponds to the quadratic selection gradient acting on a trait, and each *γ*_*ij*_ (off-diagonal) corresponds to the correlational selection gradient acting on a particular trait combination. Note that we ommitted the upper triangle of each matrix for clarity since it is simply the reflection of the lower triangle. Subtracting these two matrices results in the curvature matrix of the adaptive landscape:

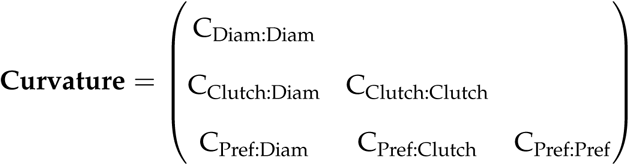

where each C_*ii*_ (diagonal) represents the effect of selection on the additive genetic variance in a trait, and each C_*ij*_ (off-diagonal) represents the effect of selection on the additive genetic covariance between a particular trait combination. We used bootstrapped values of each selection gradient to estimate the curvature of each component of the matrix and its associated 95% CI. We also used this information to determine whether the curvature of each component differed between our food-web treatments.

Knowledge of the curvature matrix alone gives an incomplete picture of its indirect effect on genetic constraints. This is because genetic constraints are ultimately determined by the structure of the G-matrix, and therefore also depend on its structure before selection. Although we do not know the underlying G-matrix for the traits we measured in this experiment, we can still gain insight to how our food-web treatment would alter genetic constraints more generally. Specifically, we calculated how our best estimate of the curvature matrix (mean values) in each treatment changed the structure of 10^4^ random G-matrices for the next generation. We restricted the range of additive genetic variance (*V*_*G*_) in these G-matrices to between 0 and 0.5 to reflect typical ranges in narrow-sense heritability values (*h*^2^; note that *h*^2^ = *V*_*G*_ when the phenotypic variance is scaled to 1). Note that this analysis assumes that the effects of recombination and mutation on the G-matrix are much smaller than the effects of selection, which appears to be a reasonable assumption over short time scales (Arnold et al., 2008).

The G-matrix itself is a complex structure, but has a clear theoretical link to the evolvability of phenotypic traits (Hansen and Houle, 2008). Evolvability measures the ability for a trait to evolve toward a given direction of selection (Hansen and Houle, 2008). In the absence of knowledge about the direction of selection that a population will actually experience in the next generation, we can compute its average evolvability over random selection gradients (Hansen and Houle, 2008; Melo et al., 2015). By computing the average evolvability (here, we used 1000 random *β*s) for each of 10^4^ random G-matrices, we can then assess how changes in the curvature matrix alter the evolutionary potential of the associated traits. We report the distribution of these effect sizes, rather than conduct a statistical test, because the number of replicates in a simulation can be arbitrarily high, thus making any effect size statistically significant (White et al., 2014).

### Adjusting for biased measurements of selection

Rather than imposing selection, parasitoids may influence the expression of herbivore traits which could bias measurements of selection. In our system, it was plausible that parasitoids may influence chamber diameter by altering larval feeding behavior or killing larvae before they complete their development. To estimate this potential bias, we subset our data to only include galls where there was variation in larval survival within the same gall (i.e. 1 > mean survival > 0). If we assume that larvae within each gall should have similar chamber diameters because they come from the same clutch and experience the same local environment (an assumption our data supports: gall ID explains 54% of the variance in chamber diameter), then the relationship between chamber diameter and larval survival in this data subset represents the effect of parasitism on trait expression (i.e. bias). We used a GLMM with the same structure as described previously except that we modeled only a linear relationship between chamber diameter and larval survival (*α*_Diam_). We detected a positive bias in both food-web treatments (original *α*_Diam_= 0.36 [0.05, 0.67]; removal *α*_Diam_= 0.42 [0.01, 0.82]), indicating that unadjusted relationships would overestimate the strength of selection on chamber diameter. To account for this bias, we subtracted our mean estimates of bias from our estimates with the full dataset prior to calculating chamber diameter’s selection surface and directional selection gradient.

### Measuring selection on egg parasitoids

Once parasitized, the gall phenotype becomes the phenotype of the egg parasitoid. This phenotype may influence the egg parasitoid’s survival in the face of larval parasitoids, and thus experiences selection. Our food-web manipulation allows us to measure selection imposed by larval parasitoids on the phenotype of egg parasitoids. Using the same models as described above, we substituted egg parasitism as our response variable to quantify selection surfaces and selection gradients acting on the egg parasitoid. Note that we cannot test the effect of food-web structure on the egg parasitoid’s adaptive landscape —we can only estimate selection imposed by larval parasitoids. This comparison is still useful though in determining the extent to which the loss of consumers may have indirect evolutionary effects by altering selection on multiple interacting populations.

All analyses and visualizations were conducted in R (R Core Team, 2018). Unless otherwise noted, we report mean estimates of selection surfaces and selection gradients with 95% confidence intervals in brackets. Note that for visualizing the adaptive landscape we restrict trait axes to ±1 SD of the mean trait value. This emphasizes the fact that we can only reliably estimate the shape of the adaptive landscape near the mean phenotype of the population (Arnold et al., 2001). We also plot mean larval survival on a natural log scale to accurately reflect the mathematical definition of the adaptive landscape (Arnold, 2003). All data and code to reproduce the reported results are publicly available on GitHub (https://github.com/mabarbour/complexity_selection) and have been archived on Zenodo (https://zenodo.org/badge/latestdoi/108833263).

## Results

### Consumer removal increases selective constraints

We found that the removal of larval parasitoids increased the number of gall midge traits under directional selection (3 of 3) relative to the original food web (1 of 3)(table 1). For example, we observed directional selection for smaller clutch sizes when we removed larval parasitoids, but there was no evidence of selection acting on this trait in the original food web (fig. 2A). This absence of selection appeared to be a result of conflicting selection pressures imposed by each guild of parasitoids. Specifically, when we subset our data to focus on differences between parasitoid guilds, we found that larval parasitoids actually impose directional selection for larger clutch sizes (larval parasitoids *β*_Clutch_= 0.13 [0.04, 0.24]). This conflicting selection is likely due to trait differences between guilds, as larger clutches may be easier targets for egg parasitoids to find, but the more complex gall structure may be more difficult for larval parasitoids to oviposit through.

**Table 1:**
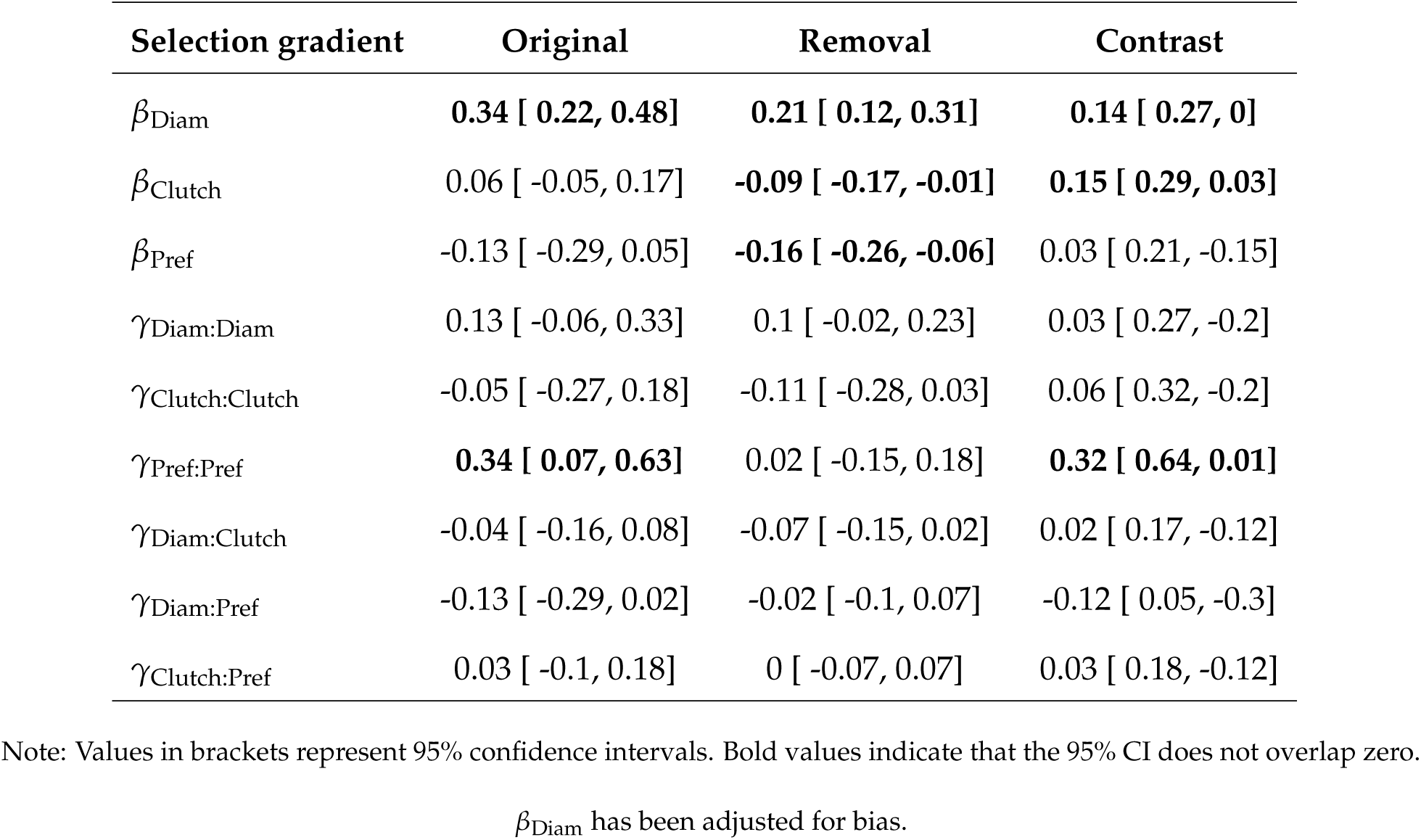
Standardized selection gradients acting on gall midges in the original food web and with the removal of larval parasitoids.

**Figure 2:**
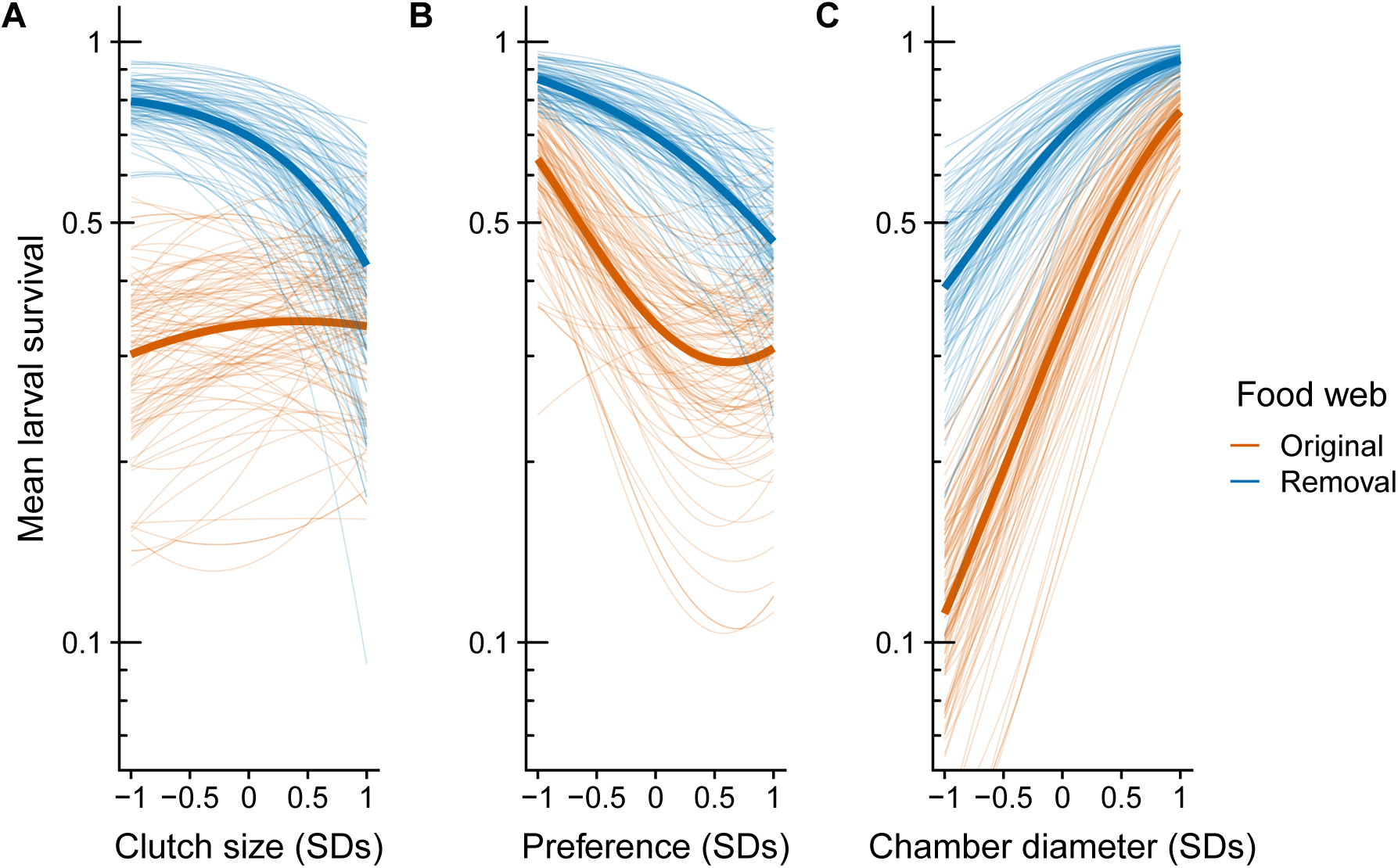
Adaptive landscape of gall midge phenotypes in the original food web and with the removal of larval parasitoids. Each panel corresponds to a different phenotypic trait: clutch size (A); oviposition preference (B); and chamber diameter (C). Bold lines represent selection experienced in the original (orange) and removal (blue) food webs. Thin lines represent bootstrapped replicates to show the uncertainty in selection. For clarity, we only display 100 bootstraps even though inferences are based on 1,000 replicates. Note that mean larval survival is plotted on a natural log scale to reflect the mathematical definition of the adaptive landscape.

We also observed clear evidence of directional selection for midges to avoid ovipositing on plants with high densities of conspecifics when we removed larval parasitoids (fig. 2B); however, the overall magnitude of selection did not differ between treatments (table 1). Still, there was no clear evidence of directional selection on oviposition in the original food web (table 1). Chamber diameter experienced positive directional selection in both food-web treatments (fig. 2C). Although the magnitude of selection on chamber diameter was relatively higher in the original food web (table 1), this was not due to any difference in the relationship between chamber diameter and survival (selection surfaces: contrast *α*_Diam_= 0.04 [0.55, −0.5]), but was simply a consequence of the (expected) lower survival of gall midges in the original food web (contrast 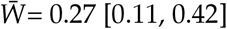). We expect this difference to be transient though, since egg parasitoids would increase in abundance once they are released from intraguild predation, which would make the strength of selection on gall diameter comparable to the original food web (if removal 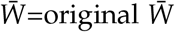, then contrast *β*_Diam_= −0.06 [-0.2, 0.1]). It is worth noting that positive selection on chamber diameter in both treatments was unexpected. For example, the fact that larger galls provide more of a refuge from larval parasitoids makes sense since they attack after the gall is formed; however, egg parasitoids attack prior to gall formation, which suggests that chamber diameter is strongly correlated with an unmeasured trait that is under selection (e.g. immune response).

To visualize the multivariate effects of these selective constraints, we plotted the adaptive landscape for each trait combination in each treatment (fig. 3). The arrows in fig. 3 represent mean estimates of directional selection gradients, while contours represent predicted survival of the mean phenotype in each food-web treatment. Notice that arrows point more toward a corner of the adaptive landscape for each combination of traits with the removal of larval parasitoids compared to the original food web. This indicates that the removal of consumers more strongly favored a specific combination of traits, rather than allowing for multiple trait combinations to have comparable fitness.

**Figure 3:**
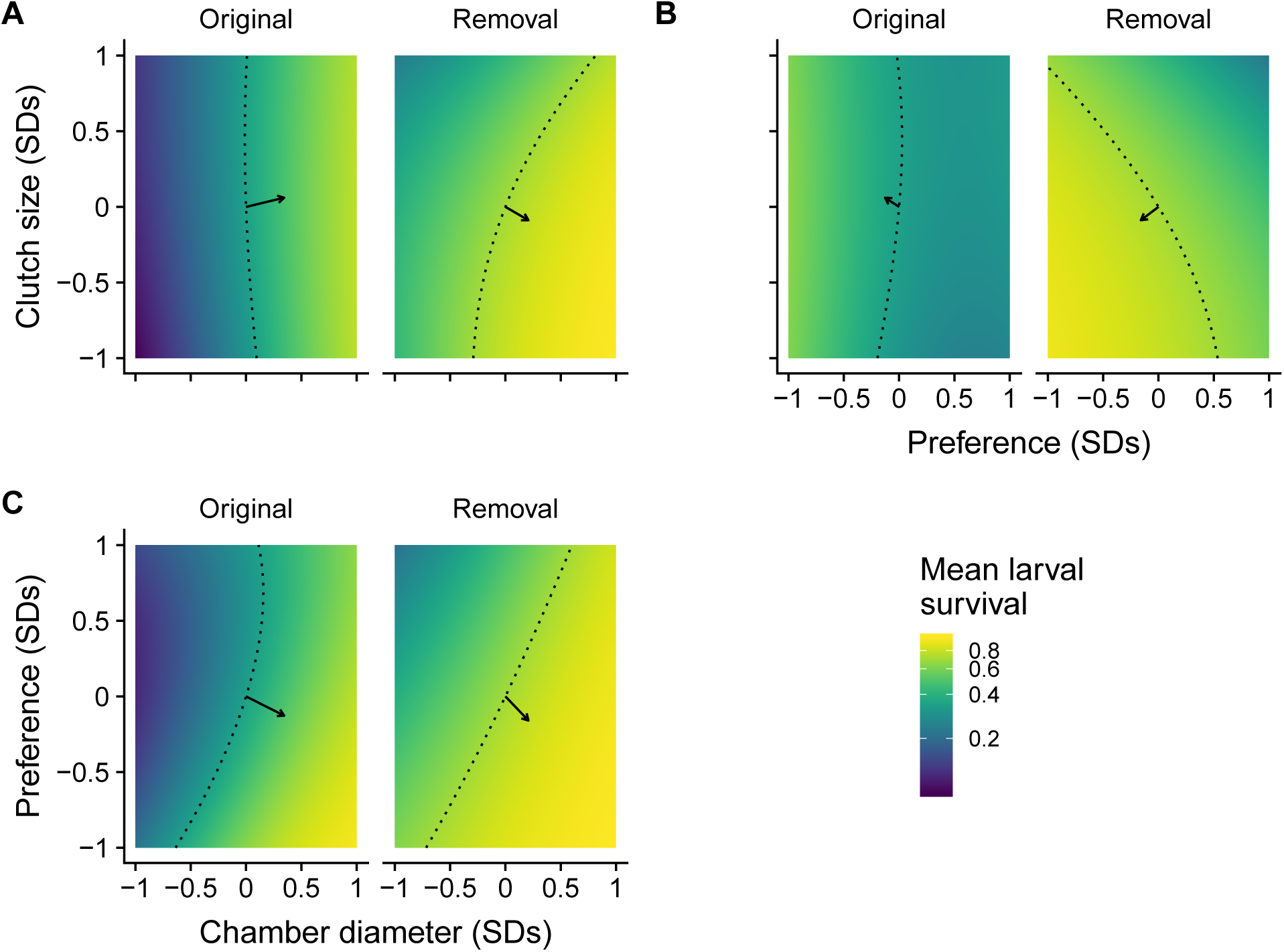
Two dimensional view of adaptive landscapes of gall midge phenotypes in the original food web and with the removal of larval parasitoids. Each panel corresponds to a different combination of phenotypic traits: clutch size and chamber diameter (A); clutch size and oviposition preference (B); oviposition preference and chamber diameter (C). Arrows represent mean estimates of directional selection gradients, while contours represent predicted larval survival of the mean phenotype in each food-web treatment. Notice that arrows point more toward a corner of the adaptive landscape for each combination of traits with the removal of larval parasitoids compared to the original food web. This indicates that the removal of consumers more strongly favored a specific combination of traits. Note that mean larval survival is plotted on a natural log scale to reflect the mathematical definition of the adaptive landscape.

### Consumer removal indirectly increases genetic constraints

The curvature of the adaptive landscape indirectly affects genetic constraints and is influenced by directional, quadratic, and correlational selection gradients. There was no clear evidence of correlational selection for any combination of traits in either food-web treatment (table 1). Similarly, there was no clear evidence of quadratic selection on chamber diameter or clutch size in either food-web treatment (table 1). In contrast, our food-web treatment did alter quadratic selection acting on oviposition preference (table 1). The negative relationship between oviposition preference and larval survival dampened at high densities in the original food web, but not with the removal of larval parasitoids (fig. 2B). This dampened relationship was partly due to a trend for nonlinear selection imposed by larval parasitoids (*γ*_Pref:Pref_= 0.18 [-0.02, 0.42]), but was also magnified by the lower average survival in the original food web. This was likely a result of larval parasitoids imposing greater mortality on egg parasitoids at high gall midge densities (see **Selection on egg parasitoids** section), and thus a less than additive effect on gall midges.

Using our estimates of directional (*β*_*i*_), quadratic (*γ*_*ii*_), and correlation selection (*γ*_*ij*_), we calculated the curvature (C = *γ* − *ββ*^T^) of the adaptive landscape in each food-web treatment.

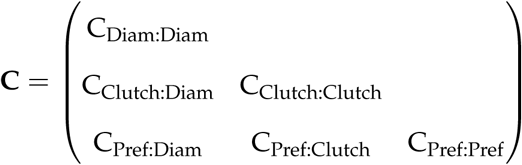

Of the different components of the curvature matrix, we found that only the curvature of oviposition preference differed between food-web treatments. Specifically, selection in the removal food web acted to decrease the additive genetic variance in preference relative to the original food web.

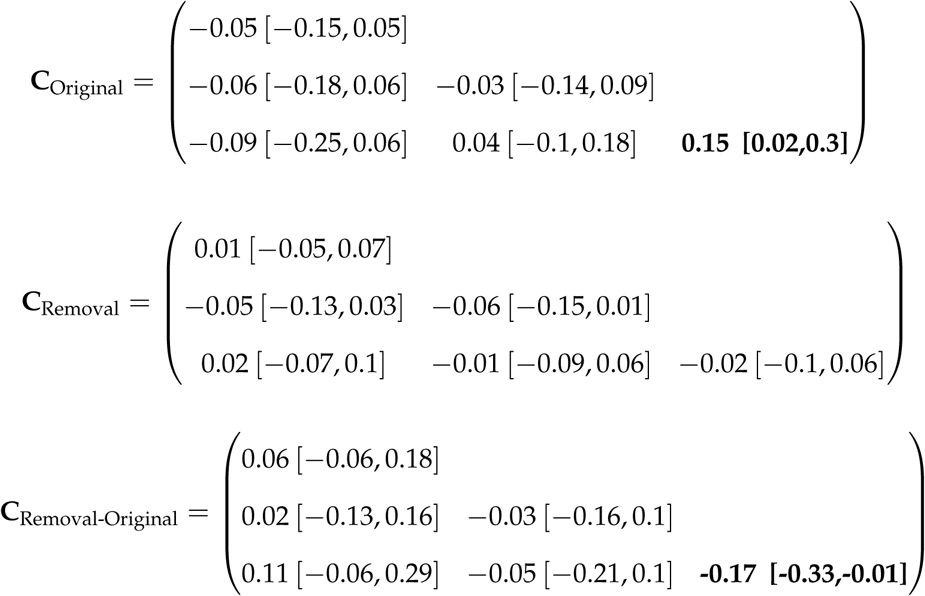

Interestingly, when we translate the effect of the curvature matrix onto genetic constraints in the next generation, we see that the removal of consumers had a general tendency to decrease evolvability (fig. 4). Specifically, the removal food web decreased the average evolvability of 71% of the random G-matrices in our simulation. If anything, we expect that this underestimates the true effect of our removal treatment. For example, if we assume egg parasitoids would eventually impose similar impact on mean fitness once they are released from intraguild predation (i.e. removal 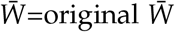), then the removal food web decreases the average evolvability in 78% of the G-matrix scenarios.

**Figure 4:**
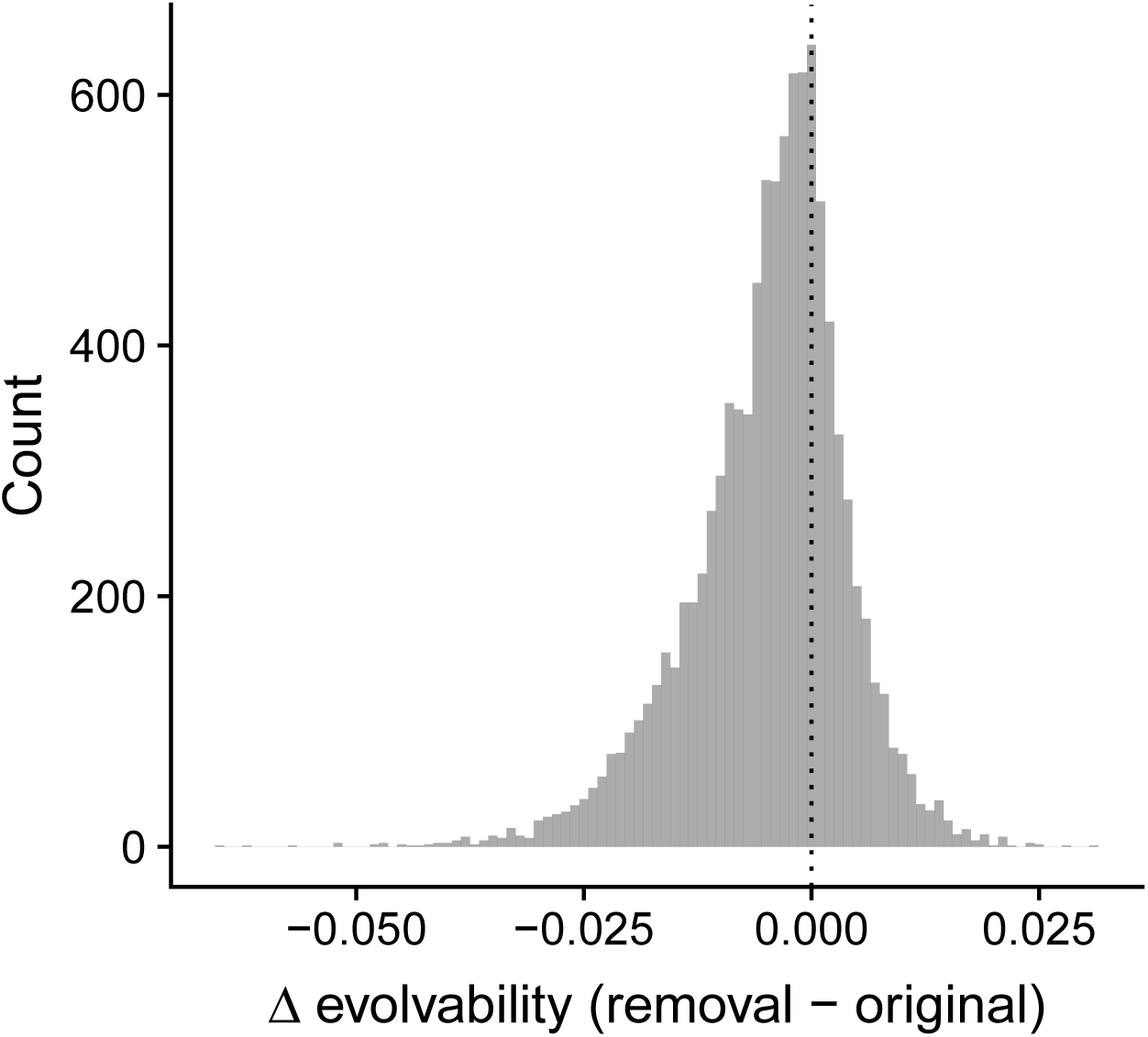
Change in average evolvability for 10,000 random G-matrices using our best (mean) estimate of the curvature matrix for each food-web treatment. We found that the curvature of the removal food web decreased evolvability in 71% of the G-matrices (i.e. the change in evolvability was negative for 71% of the simulations), suggesting that the removal of consumers tended to decrease evolutionary potential of traits in our study.

### Selection on egg parasitoids

The removal of larval parasitoids not only had direct effects on gall midge fitness, but also imposed indirect effects that would be felt in the next generation. For example, the removal of larval parasitoids altered the relationship between gall midge preference and the probability of observing egg parasitoids (*γ*_Pref:Pref_=-0.46 [−1.07, −0.02], table S1), such that the impact of larval parasitoids increased nonlinearly with higher gall midge densities (fig. 5).

**Figure 5:**
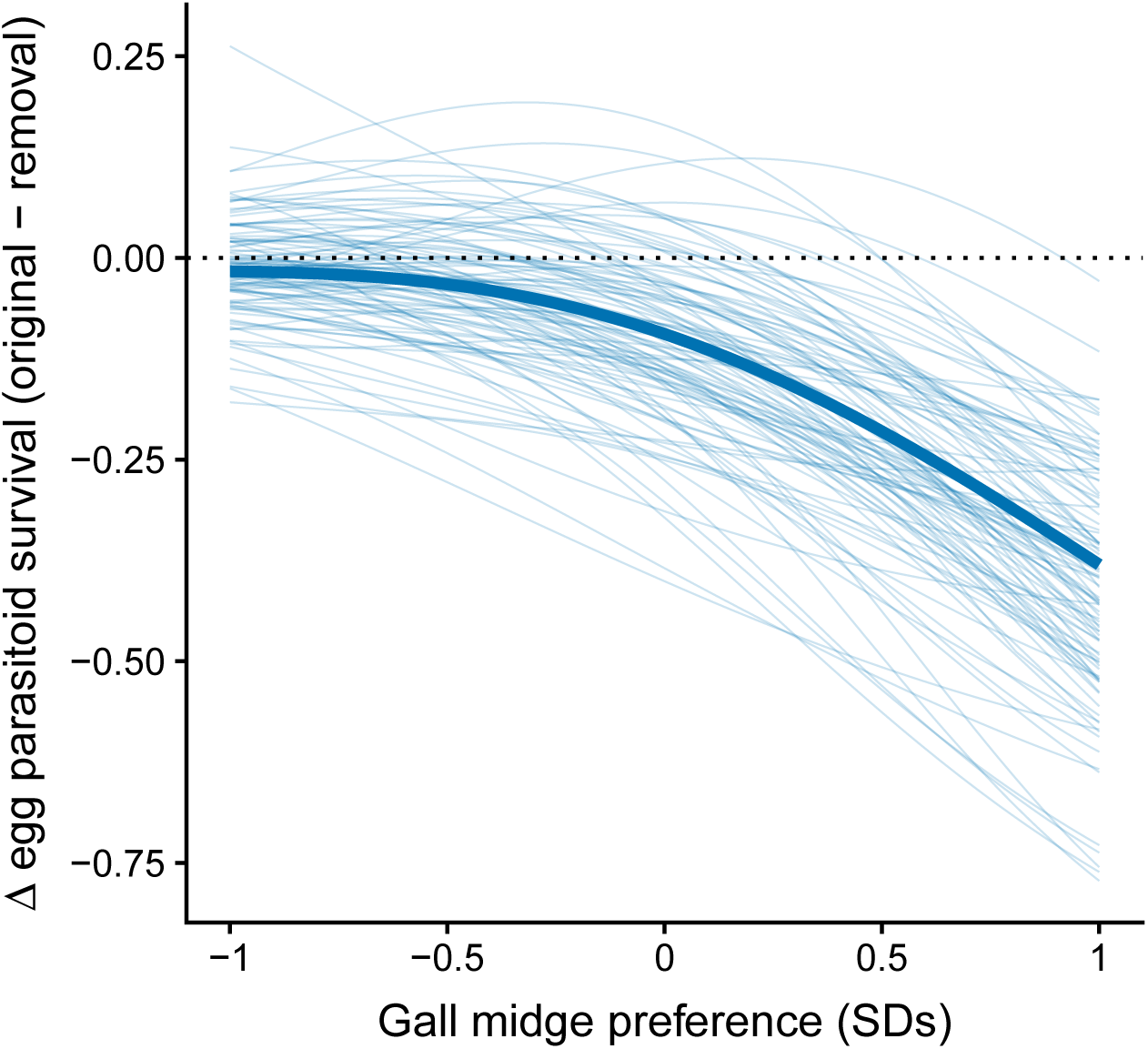
Selection imposed by larval parasitoids on egg parasitoids (*Platygaster* sp.). The bold line represents the average difference in the probability of observing the egg parasitoid (original minus removal of larval parastioids) as a function of gall midge oviposition preference. Thin lines represent bootstrapped replicates to show the uncertainty in selection. For clarity, we only display 100 bootstraps even though inferences are based on 1,000 replicates. The decrease in the probability of observing egg parasitoids at high gall-midge densities indicate that larval parasitoids impose nonlinear selection on egg parasitoids.

## Discussion

We found that the removal of larval parasitoids constrained phenotypic evolution in gall midges in two key ways. First, more traits contributed to the slope of the adaptive landscape in the absence of larval parasitoids, suggesting greater constraints on the trajectory of phenotypic evolution. Second, removing larval parasitoids altered the curvature of the adaptive landscape in such a way that tended to decrease the evolvability of associated traits. Assuming these traits have other ecological functions, then this decrease in evolvability could constrain the gall midge’s adaptive potential in the face of novel selection pressures. Our experiment also revealed evidence of indirect selection pressures, suggesting that the loss of consumers may have complex effects on the trajectories of phenotypic evolution. Taken together, our study provides experimental evidence from the field that the loss of consumers may constrain the adaptive potential of remaining populations.

The generality of our results likely depends on the relative abundance and functional differences between consumers in a community. For example, if consumers do not differ from each other, then we do not expect the loss of consumers to modify selective constraints. Also, many consumers may be at too low of abundances to impose selection on their resources. Rank abundance curves (Preston, 1948) and the disproportionate number of weak interactions in diverse communities (Paine, 1992) support this notion. This logic suggests that we may not have observed the effects we did if we had only removed one larval parasitoid, because each species had relatively low abundance (Barbour et al., 2016) and they likely share similar ecological roles. When consumers are functionally different and abundant though, the effect of consumer loss will depend on whether different consumers impose conflicting selection pressures or select for distinct traits. When consumers impose conflicting selection on traits, as in our study and others (Weis and Abrahamson, 1985; Abrahamson and Weis, 1997; Start and Gilbert, 2016; Start et al., 2018), then consumer diversity acts to neutralize selection and relax selective constraints. On the other hand, different consumers may impose selection on different traits; therefore, a more diverse consumer community may favor a particular combination of traits and increase selective constraints. Examples of this include strong genetic covariances in plant resistance to different insect herbivores (Maddox and Root, 1990; Wise, 2007; Wise and Rausher, 2013), although there are also examples where these covariances are weak (Roche and Fritz, 1997; Barbour et al., 2015), or vary from year-to-year (Johnson and Agrawal, 2007). We suggest that gaining predictive insight to the evolutionary consequences of food-web disassembly requires an understanding of the mechanisms governing the assembly of trophic interactions (Bascompte and Stouffer, 2009).

We also found evidence for a general decrease in trait evolvability when we excluded larval parasitoids due to changes in the curvature of the adaptive landscape. This result was driven by nonlinear selection on gall midge oviposition preference in the original food web, which was due to both increases in intraguild predation and the lower mean survival of gall midges. While these results are intriguing, they still indicate that the indirect effects of selection are contingent on the structure of a population’s G-matrix (e.g. consumer removals *increased* evolvability in 29% of the scenarios). This highlights the importance of future studies to characterize the G-matrix in order to accurately predict how changes in network context will alter the evolutionary potential of remaining populations. Indeed, current theory often assumes genetic variances and covariances remain constant over time rather than dynamically changing with the network context (McPeek, 2017; Guimarães et al., 2017). Our empirical results highlight the need to explore the evolutionary consequences of not only direct effects of selection, but indirect effects on genetic constraints that emerge in a network of interacting species.

An important caveat of our study is that we did not do a factorial manipulation of both parasitoid guilds, making it difficult to conclude whether our results would change if we manipulated the presence/absence of the dominant egg parasitoid. If we assume that higher-order interactions (Levine et al., 2017) are weak between parasitoid guilds, then we can gain insight to how the loss of the egg parasitoid would alter selection by isolating the contribution of larval parasitoids to selection in our original food-web treatment. When we do this, we see the same qualitative effects as we do when we removed larval parasitoids. For example, we see clear evidence of all three traits being under directional selection (i.e. greater selective constraints, table S2) as well as a decrease, albeit smaller, in trait evolvability under different G-matrix scenarios (57%, fig. S1). This suggests that our results could be robust to this caveat, which was simply not possible to manipulate given the biology of our system (see **Manipulating Food-web Structure** section for explanation).

Our results suggest that the loss of consumers may not only directly affect connected species, but also result in indirect evolutionary effects. In our study, this indirect effect arises from egg parasitoids being released from intraguild predation when we excluded larval parasitoids. This release occurs more on trees with high larval densities, which could intensify future selection on gall midge oviposition preference. A growing number of experiments over the past two decades have demonstrated the presence and potential importance of indirect evolutionary effects that emerge in ecological communities (Pilson, 1996; Juenger and Bergelson, 1998; Stinchcombe and Rausher, 2001; Lankau and Strauss, 2007; Walsh and Reznick, 2008, 2010; terHorst, 2010; Sahli and Conner, 2011; Lau, 2012; terHorst et al., 2015; Schiestl et al., 2018; Start et al., 2019). If indirect evolutionary effects are common (Miller and Travis, 1996; Walsh, 2013; Guimarães et al., 2017), then predicting evolutionary trajectories resulting from the loss of consumers will require evolutionary studies to explicitly account for the ecological networks that species are embedded in.

Our study gives insight to how the loss of consumers alters evolutionary constraints on remaining populations. In particular, it hints at a potential insidious effect of local extinctions that compromises the robustness of remaining populations to future environmental change. Our work also highlights some key challenges for predicting phenotypic evolution in rapidly changing communities. For example, many theoretical models of eco-evolutionary dynamics focus on phenotypic change in a single trait, yet our results highlight that the number of traits under selection may change with the network context. Importantly, we found that different species/guilds imposed different selection pressures. Knowing these hidden selection pressures is critical for prediction, because the trajectory of evolution will depend on the nature of change in the ecological community. We expect that a continued integration of adaptive landscapes and ecological networks will enhance our ability to predict the evolutionary consequences of changes in ecological communities.

## Acknowledgements

We thank the staff of Humboldt Bay National Wildlife Refuge (U.S. Fish and Wildlife Service) for facilitating experimental logistics. For assistance with fieldwork, we thank Ruthie Espanol and Andrew MacDonald. For funding support, we thank the University of British Columbia (James Robert Thompson Fellowship and Four-Year Fellowship to M.A. Barbour), NSERC (Discovery grant to Greg Crutsinger), and the Swiss National Science Foundation (grant 31003A_160671 to J. Bascompte). The authors declare no conflict of interest.

## Author Contributions

M.A.B. conceived the idea behind the study and designed the field experiment. M.A.B. and B.L. setup and conducted the experiment. M.A.B., A.S., and C.J.G. collected the data. M.A.B. analyzed the data. M.A.B. wrote the manuscript with primary input from J.B. and additional feedback from C.J.G.

## Data Accessibility

All data and code to reproduce the reported results are publicly available on GitHub (https://github.com/mabarbour/complexity_selection) and have been archived on Zenodo (https://zenodo.org/badge/latestdoi/108833263).

## Notes

https://zenodo.org/badge/latestdoi/108833263

