## Supplementary Material for "Loss of consumers constrains phenotypic evolution in the resulting food web"

Table S1: Standardized selection gradients acting on egg parasitoids (*Platygaster* sp.)

| **Selection gradient** | **Contrast = Original - Removal** |
|---|---|
| $\beta_{\text{Diam}}$ | -0.03 [ -0.3, 0.25] |
| $\beta_{\text{Clutch}}$ | 0.07 [ -0.26, 0.39] |
| $\beta_{\text{Pref}}$ | -0.25 [ -0.64, 0.09] |
| $\gamma_{\text{Diam:Diam}}$ | -0.05 [ -0.43, 0.33] |
| $\gamma_{\text{Clutch:Clutch}}$ | -0.21 [ -0.68, 0.26] |
| $\gamma_{\text{Pref:Pref}}$ | **-0.46 [ -1.07, -0.02]** |
| $\gamma_{\text{Diam:Clutch}}$ | 0 [ -0.29, 0.27] |
| $\gamma_{\text{Diam:Pref}}$ | 0.25 [ -0.04, 0.6] |
| $\gamma_{\text{Clutch:Pref}}$ | -0.18 [ -0.52, 0.12] |

Note: Values in brackets represent 95% confidence intervals. Bold values indicate that the 95% CI does not overlap zero.

$\beta_{\text{Diam}}$ has been adjusted for bias.

Table S2: Standardized selection gradients imposed by larval parasitoids on gall midges in the original food web. This gives insight to selection acting on gall midges in the absence of the dominant egg parasitoid.

| **Selection gradient** | **Larval Parasitoids** |
|---|---|
| $\beta_{\text{Diam}}$ | **0.23 [ 0.13, 0.36]** |
| $\beta_{\text{Clutch}}$ | **0.13 [ 0.04, 0.24]** |
| $\beta_{\text{Pref}}$ | **-0.17 [ -0.34, -0.03]** |
| $\gamma_{\text{Diam:Diam}}$ | 0.06 [ -0.07, 0.2] |
| $\gamma_{\text{Clutch:Clutch}}$ | 0.01 [ -0.14, 0.15] |
| $\gamma_{\text{Pref:Pref}}$ | 0.18 [ -0.02, 0.42] |
| $\gamma_{\text{Diam:Clutch}}$ | 0.02 [ -0.07, 0.11] |
| $\gamma_{\text{Diam:Pref}}$ | -0.05 [ -0.18, 0.06] |
| $\gamma_{\text{Clutch:Pref}}$ | -0.04 [ -0.14, 0.06] |

Note: Values in brackets represent 95% confidence intervals. Bold values indicate that the 95% CI does not overlap zero. $\beta_{\text{Diam}}$ has been adjusted for bias.

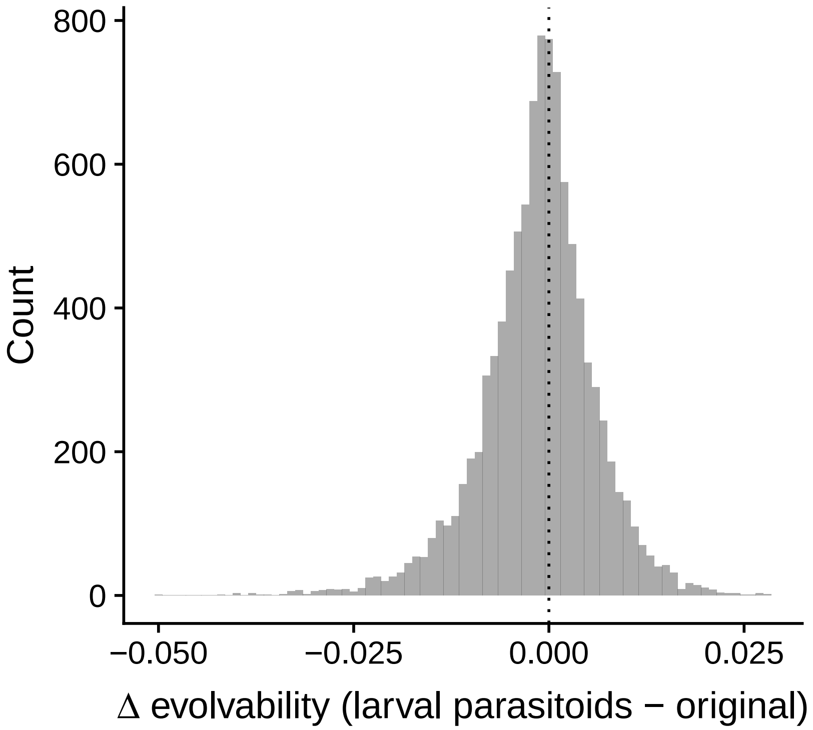


Figure S1: Change in average evolvability for 10,000 random G-matrices using our best (mean) estimate of the curvature matrix for selection in the absence of egg parasitoids vs. the original food web. We found that the curvature imposed by the simulated removal of egg parasitoids decreased evolvability in 57% of the G-matrices (i.e. the change in evolvability was negative for 57% of the simulations), which is smaller in magnitude, but in the same direction, as the effects of removing larval parasitoids.
